# Highly parallel direct RNA sequencing on an array of nanopores

**DOI:** 10.1101/068809

**Authors:** Daniel R. Garalde, Elizabeth A. Snell, Daniel Jachimowicz, Andrew J. Heron, Mark Bruce, Joseph Lloyd, Anthony Warland, Nadia Pantic, Tigist Admassu, Jonah Ciccone, Sabrina Serra, Jemma Keenan, Samuel Martin, Luke McNeill, Jayne Wallace, Lakmal Jayasinghe, Chris Wright, Javier Blasco, Botond Sipos, Stephen Young, Sissel Juul, James Clarke, Daniel J Turner

## Abstract

Ribonucleic acid sequencing can allow us to monitor the RNAs present in a sample. This enables us to detect the presence and nucleotide sequence of viruses, or to build a picture of how active transcriptional processes are changing – information that is useful for understanding the status and function of a sample. Oxford Nanopore Technologies’ sequencing technology is capable of electronically analysing a sample’s DNA directly, and in real-time. In this manuscript we demonstrate the ability of an array of nanopores to sequence RNA directly, and we apply it to a range of biological situations. Nanopore technology is the only available sequencing technology that can sequence RNA directly, rather than depending on reverse transcription and PCR. There are several potential advantages of this approach over other RNA-seq strategies, including the absence of amplification and reverse transcription biases, the ability to detect nucleotide analogues and the ability to generate full-length, strand-specific RNA sequences. Direct RNA sequencing is a completely new way of analysing the sequence of RNA samples and it will improve the ease and speed of RNA analysis, while yielding richer biological information.

## Introduction

The overall aims of transcriptomics include comprehensive sequencing of all species of transcript in a sample, determining the transcriptional structure of genes including splice variants and fusion genes, quantifying changing expression levels of transcripts in different developmental stages and in different environments, and ascertaining the level of antisense transcription occurring in an organism’s cells^1^.

To fully satisfy these aims, a method of detecting and characterizing RNA is required that is accurate, strand-specific, quantitative across a wide dynamic range, does not need prior knowledge of the genome sequence or the transcripts that are being assayed, is capable of revealing the presence and identity of modified bases, and can detect antisense transcripts without any possibility that these are library preparation artefacts^2^. It would also be advantageous to be able to generate sequence reads that are sufficiently long to span multiple splice junctions, for unambiguous mapping. There is no commercially-available technology which meets all of these requirements.

Transcriptomic analyses have been made possible by the massive throughput of next-generation sequencing-by-synthesis platforms: sequencing of complementary DNA (termed RNA-seq) has enabled us to build an accurate picture of the active transcriptional patterns within organisms^1^. A single run on such a sequencer can generate millions of individual short reads,providing a large dynamic range with no upper limit for quantification, potentially allowing precise quantification of all transcripts without any prior knowledge of their sequence, or even their existence.

The most commonly used strategy for next generation RNA-seq involves either polydeoxythymine (poly dT) priming or fragmentation of ribonucleic acid (RNA) followed by complementary DNA (cDNA) synthesis with random hexamer priming. These cDNA strands are then typically prepared for analysis on a given sequencing system using a library preparation method that incorporates PCR. However, wherever PCR is used in a library preparation, bias is introduced^3^. For example, the resulting library will have reduced complexity compared to the total mRNA pool, because not all transcripts will amplify with the same efficiency, causing drop-out of some RNA species, and excessive amplification of other species. Such PCR duplicates are difficult to distinguish from genuinely abundant RNA species. Additionally, PCR amplification creates a synthetic copy of the original RNA strand, and any epigenetic information is lost. Because of this, it would be an advantage to avoid amplification in the library prep altogether^4^.

Exceptions to PCR-based library preparations include FRT-seq on the Illumina platforms, in which the first strand cDNA synthesis reaction is performed on single strands of RNA which are hybridised to the flowcell surface^5^, and the Direct RNA sequencing on the Helicos platform^6^, where a sequencing-by-synthesis reaction was performed using native RNA strands as the sequencing template. In both of these cases, PCR amplification is avoided, removing this source of bias, and meaning that the data obtained should reflect the abundance of RNAs in the original sample more faithfully. However, these approaches still rely on synthetic copies of the original RNA strand, so do not retain information about modifications. Both of these approaches necessarily generate short sequence reads, which is far from ideal because in eukaryotes, transcripts commonly undergo alternative splicing, which creates multiple different isoforms^7^. Gene isoforms can have different transcription start sites, coding sequences and untranslated regions, in some cases producing very different functions for the different isoforms^8^. Short read sequencing typically cannot span entire transcripts and frequently does not span both sides of splice junctions adequately, and in either case new splice variants can be missed^9^.

By priming cDNA synthesis from the poly-A tail of eukaryotic transcripts and following a strand-switching protocol, it is possible to create a library consisting almost entirely of full-length cDNA strands. The power of coupling this preparation method with a long read sequencing platform has been shown recently^10,11^. In both cases, the authors were able to obtain full gene structures without the need for assembly, and discovered a large number of new transcript isoforms. The combination of a strand-switching protocol with a long read sequencer allows generation of strand-specific data, which is essential for comprehensive annotation of the transcriptome^12-14^ and to identify antisense transcription, a mechanism for gene regulation and gene silencing that has been reported to be widespread in the mammalian transcriptome^15,16^

In none of these methods, even the ‘Direct RNA’ approach^6^ is the RNA strand from the organism being analysed directly; the sequencing reactions are based on the detection of the products of a synthesis reaction. This means that library preparation is subject to the processivity and error-rate limitations of reverse transcription. Even if a strand-switching protocol is followed, which selects only those cDNA strands that correspond to the full length of the original transcript, poor reverse transcriptase processivity means that longer transcripts, or those which are difficult to reverse transcribe, are less well represented than shorter ones. This bias is exacerbated during PCR amplification. In addition, because reverse transcriptases are capable of using either RNA or DNA as a template, there is the opportunity to create chimeric products and other artefacts^2^. Furthermore, because cDNA-based approaches to RNA-seq detect enzymatically-incorporated nucleotides rather than the original RNA strand that was in the sample of interest, it is not possible to detect base modifications directly.

Oxford Nanopore Technologies have developed a nanopore-based sensing platform which is capable of sequencing DNA directly without the need to perform an enzymatic synthesis reaction. As with DNA oligonucleotides, strands of RNA can be made to pass through a protein nanopore embedded in a hydrophobic membrane when an electrical potential is applied^17^ If a motor protein is used to control the speed of translocation of the RNA strand through the nanopore, a signal can be obtained which is governed by the identity of the RNA strand. The signal changes if the primary sequence of the RNA is changed^18^.

Here we demonstrate that our sensing platform is capable of sequencing RNA and RNA modifications directly, without amplification. Oxford Nanopore’s MinION uses an array of single protein nanopores, each of which is embedded in a synthetic polymer membrane. At either side of this membrane is a salt-containing buffer, and when an electrical potential is applied, ions pass through the nanopores. We are able to measure the current flowing through the individual nanopores, and when a strand of DNA or RNA passes through a pore, the flow of current is partially blocked, in a sequence-specific way. We limit the speed with which the strands pass through the pore using a motor protein which is attached to the template strands during library preparation, but which does not process along the strand until the strand is captured by a nanopore. Using a custom Hidden Markov Model or Recurrent Neural Network we are able to convert the current readings to basecalled sequences.

There are several potential advantages to direct RNA sequencing with nanopores: i) the approach does not require any prior knowledge of the sequences being detected, and is thus capable of detecting unknown transcripts; ii) it is necessarily strand-specific; iii) it detects the native nucleotides, not products of a synthesis reaction, meaning that all nucleotides in the strand, including those that are modified, synthetic, or irregular have the potential to be identified; iv) by definition, PCR and other forms of amplification are not used, which removes all amplification biases; v) the system is capable of sequencing strands that are tens of kilobases in length, meaning that given a fully processive motor protein, the only limitation on read length is the sample quality; and vi) it is not affected by the processivity and error-rate limitations of reverse transcription.

This is, to our knowledge, the first parallel truly direct RNA sequencing method. The method is performed using a consumable chip on a commercially available handheld device, and is compatible with real-time data analysis.

## Methods

### i. Adapter preparation

#### a. Double-stranded adapter hybridisation (RT splint adapter and sequencing adapter)

A 4uM stock of hybridized ‘RT splint’ adapter was prepared as follows:

- 2 µl of 100 µM top strand adapter
- 2 µl of 100 µM bottom strand adapter
- 1 µl of 50x annealing buffer (0.5 M Tris.HCl pH 7.5, 2.5 M NaCl)
- 45 µl of deionized water

The mixture was heated to 95 °C for 2 minutes and was then cooled to 20 °C at a rate of 0.5 °C sec^-1^

#### b. Pre-loaded sequencing adapter

The motor protein was buffer exchanged into a buffer consisting of 100 mM potassium phosphate pH 8, 125 mM NaCl, 5 mM EDTA, 0.1% TWEEN-20, and then incubated for 10 minutes with 1 nM hybridised sequencing adapter, to allow binding. The adapter / enzyme complex was purified using a 3.7x excess of AMPure beads before elution into 50 mM Tris pH 8, 20 mM NaCl. We assume a concentration of 100 nM.

### ii. Library preparation

We prepared direct RNA libraries using the library prep depicted in Fig. 1. Briefly, 500 ng of poly A+ yeast RNA was ligated to 100 nM pre-annealed RT splint adapter using T4 DNA ligase. The products were purified using a 1.8 μl of of Agencourt AMPure beads per μl of sample, washing twice in 70% ethanol. A reverse transcription reaction was performed on the products, using the RT splint as a primer, before a second 1.8x AMPure purification. Sequencing adapters preloaded with motor protein were then ligated onto the cDNA: mRNA hybrid duplex, the tether oligonucleotide was added, and the final library was cleaned up using 1.8 μl of of AMPure beads per μl of sample. The library was then mixed with running buffer to 150 μl and injected into an R9 flow cell and run on a Mk1b MinION.

**Fig. 1.**
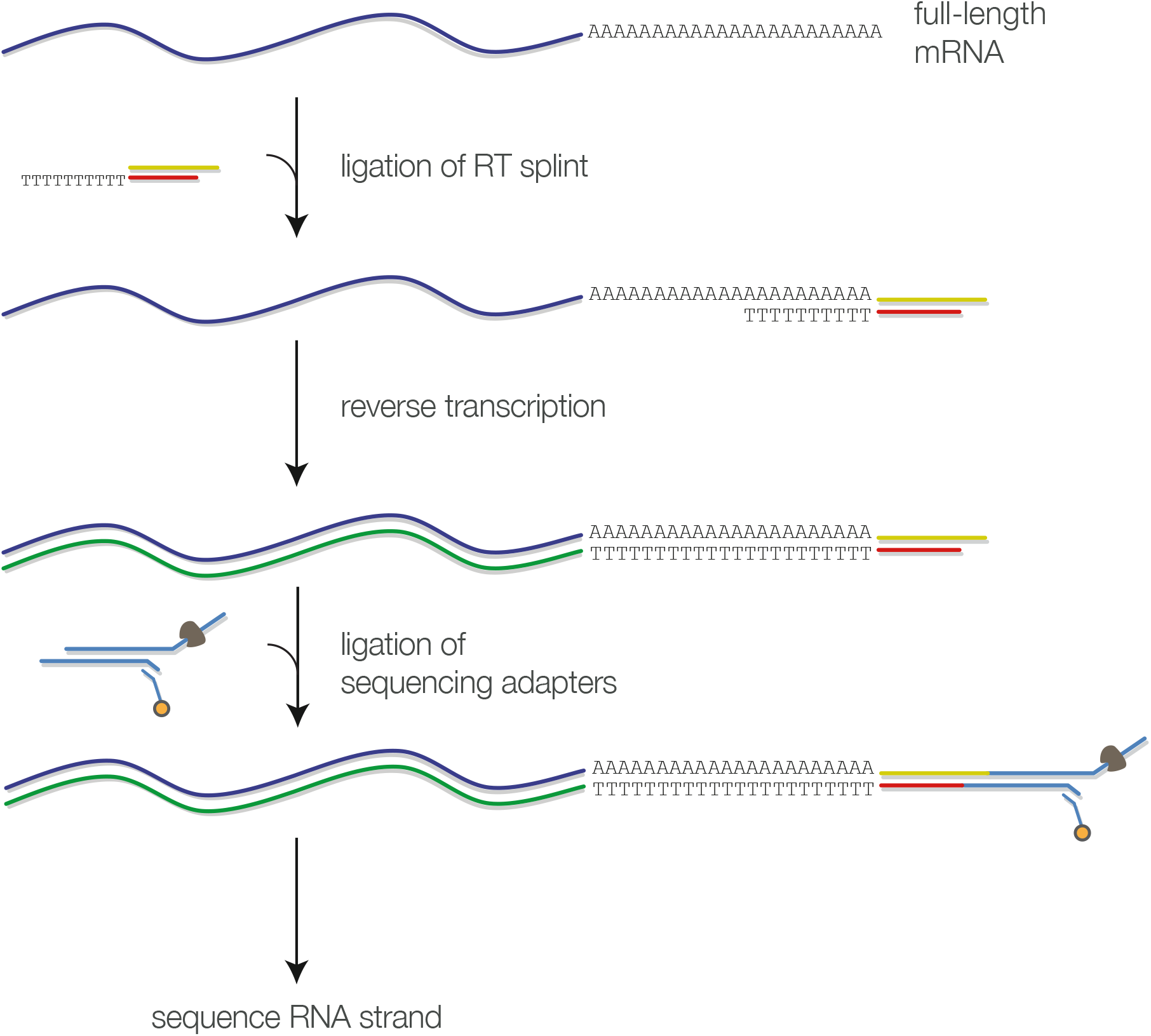
One of several possible library preparation methods for Direct RNA sequencing.

### iii. Data analysis

#### a. Basecalling

To decode nucleotide sequences from our Nanopore ionic current measurements we employ Hidden Markov Models (HMMs). We assume that the ionic current depends solely on five nucleobases resident in the nanopore, giving us an HMM with 1024 states, or kmers. The stepwise nature of the ionic current events is thereby explained as observations of the discrete kmers; an event being a grouping of raw data points with mean value μ. For each event the value of μ is assumed to be drawn from a normal distribution with mean μ_k and standard deviation σ_k, the index k identifying a specific kmer. These emission parameters have been trained using standard methods with data collected from several MinION runs using a range of sequences. Our HMM is constrained by allowing transitions primarily between the hidden states which correspond to 0, 1, or 2 nucleobase movements (often referred to as stay, step, and skip). We allow, by a small uniform probability, transitions from the current state to any other state. With the HMM thus specified, kmer sequences are decoded with an iterative variation of the posterior-Viterbi decoding scheme^19^. During iterations, scaling parameters and updates to the HMM transition matrix (the stay, step, and skip probabilities) are made^20^. To finally provide a basecall, the kmer predications are joined in a maximally overlapping manner.

#### b. Human rhinovirus

Basecalls for a Human Rhinovirus (HRV) dataset were produced using the HMM described above. Alignments to the reference were made using LAST^21^ parameterised –a 50 –b 100 –e 1200 –y 7000, and using a custom mismatch matrix derived previously. Alignments were filtered to those covering at least 60% of the read.

#### c. Read mapping and counting

To map reads to the Saccharaomyces cerevisiae Ensembl release 85 cDNA collection we performed basecalling with the HMM described above and used bwa mem^22^ with the parameters –W 15 –k 8 –x ont2d. Pairwise alignments were created with Clustal W^23^. To quantify the synthetic Lambda phage transcripts we performed basecalling as before and performed alignments using LAST as with the HRV samples. For each read the highest scoring alignment was used to identify the origin of the RNA molecule.

#### d. Detection of methylated adenosine

To show that nanopore sequencing can detect the presence of methylation we trained HMMs from two distinct samples: a sample which contains only canonical A bases and a sample which exclusively contains m6A bases. For simplicity of exposition we choose to model all positions of the reference sequence independently, that is, the HMM comprises a state-space containing as many states as there are bases in the reference sequence. Such modelling allows us to relax the assumption that only five bases contribute to the observed ionic current. To bootstrap the training we do however use the emission parameterisation of our 5mer basecalling model: we initialise the parameters μ_p and σ_p to those which can be found in our 5mer model indexed on the 5 bases surrounding the reference position p. Having trained the HMM models, the final emission parameter sets represent a consensus ^“^squiggle^”^ across all reads in the two datasets. To compare the consensuses we perform least squares regression of μ_p and μ_p(meth) at reference positions which are expected to not be affected by the methylation under the 5mer assumption.

## Results

### i) Direct RNA sequencing of yeast transcripts

When an RNA strand passes through a nanopore, the current level changes with the sequence in a similar way to when the analyte is DNA. We prepared a 1D RNA library from yeast poly A+ RNA, following the library preparation protocol outlined in **Fig. 1** and sequenced the library using R9 flowcells, at a translocation speed of ˜90 nucleotides per second per pore on a MinION. We measured current levels using and created fast5 files for each read. **Fig. 2a** shows the current levels resulting from the translocation of a single RNA molecule through one of the pores in the array. When no strand is passing through a given pore, then the maximum ionic current is able to flow, causing an ‘open pore current’. As a strand begins to pass through the pore in a 3’-5’ direction, the adapter oligo is detected first, followed by the poly A tail of the transcript, then the body of the transcript, and the current returns to the open pore level as the transcript exits the pore on the opposite side of the membrane.

**Fig. 2.**
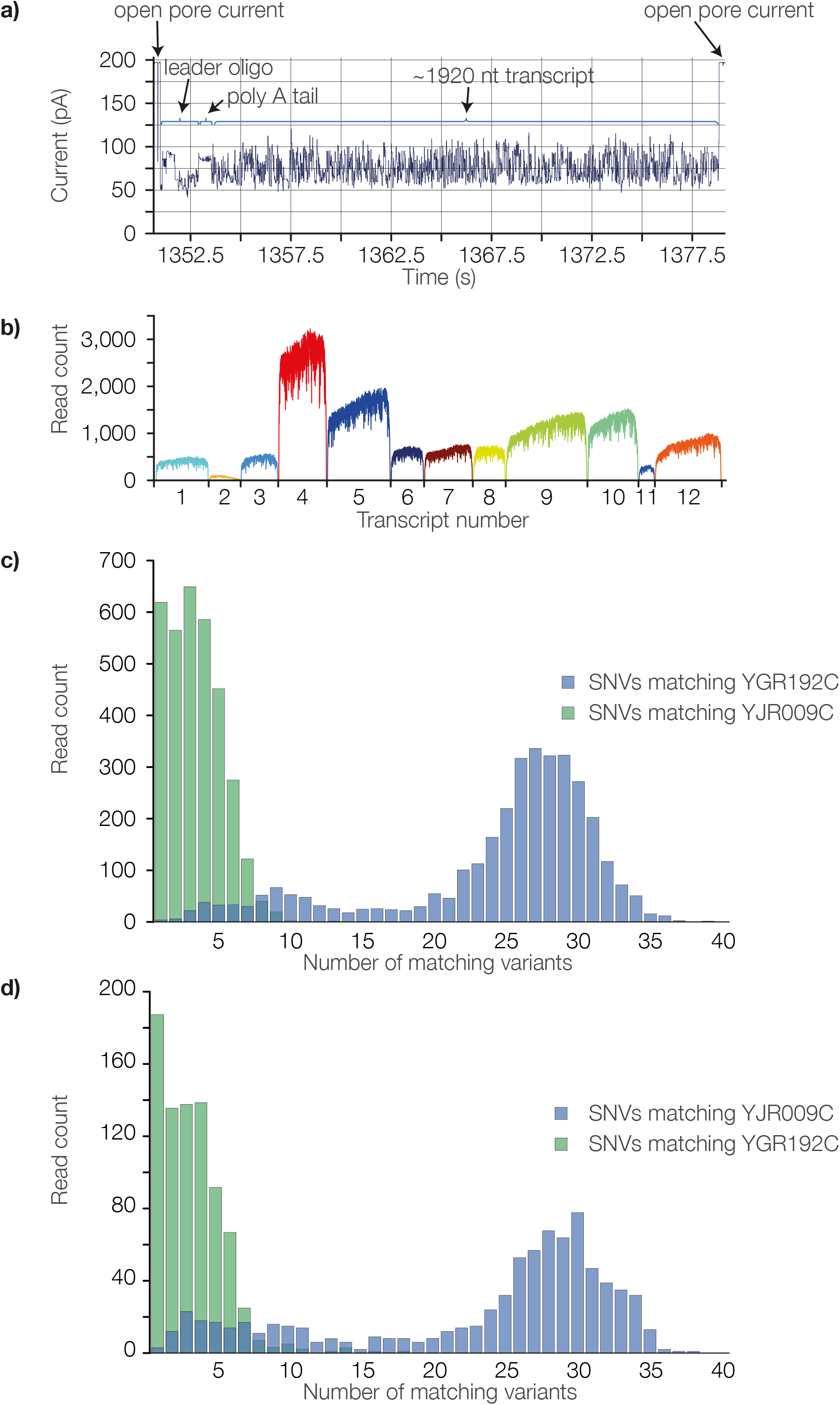
Direct RNA sequencing of yeast poly A+ RNA. **a)** Squiggle resulting from translocation of a single nscript through a pore in the array. The trace is annotated with significant features **b)** Nucleotide coverage for nscripts in the sample identified overa range of read counts **c)** Variants matching each isozyme reference quence for all reads mapping to YGR192C **d)** Variants matching each isozyme reference sequence for all read mapping to YJR009C

Fast5 files were basecalled and aligned to the Saccharomyces cerevisiae S228C transcriptome. **Table 1** shows an alignment of a section of a typical read against the closest match in the transcriptome, YDL229W. We estimate the identity of the complete read to be ˜80%.

**Table 1.**
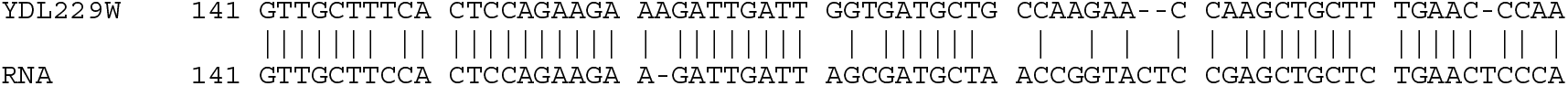
Alignment of Direct RNA sequence data to the closest match from the S228C transcriptome

**Fig. 2b** and **Table 2** show a selection of abundant poly A+ RNAs identified from the yeast transcriptome, along with their identities. As might be expected for a growing yeast culture, these transcripts are mostly ribosomal or involved in metabolism. The coverage of several of these transcripts appears to decrease towards the 5’ end, raising the possibilities that further engineering of the current RNA motor protein may improve its processivity, and that more careful handling of RNA strands may help to reduce degradation.

**Table 2.** Identities and locations of the most abundant transcripts identified

|  |  |  |  |  |  |  |  |
| --- | --- | --- | --- | --- | --- | --- | --- |
| 1 | YBR031W | II | 300166 | 301254 | 1,088 | 527 | Ribosomal 60S subunit protein L4A |
| 2 | YCR013C | III | 138402 | 139049 | 647 | 108 | Unknown |
| 3 | YGL123W | VII | 277617 | 278381 | 764 | 574 | Protein component of the small (40S) ribosomal subunit |
| 4 | YGR192C | VII | 882812 | 883810 | 998 | 3,217 | Glyceraldehyde-3-phosphate dehydrogenase (GAPDH), isozyme 3 |
| 5 | YHR174W | VIII | 451327 | 452640 | 1,313 | 2,029 | Enolase II, a phosphopyruvate hydratase |
| 6 | YJR009C | X | 453683 | 454681 | 998 | 743 | Glyceraldehyde-3-phosphate dehydrogenase (GAPDH), isozyme 2 |
| 7 | YJR123W | X | 651901 | 652578 | 677 | 754 | Protein component of the small (40S) ribosomal subunit |
| 8 | YLR044C | XII | 232390 | 234081 | 1,691 | 1,514 | Major pyruvate decarboxylase isozyme |
| 9 | YLR075W | XII | 282927 | 283592 | 665 | 720 | Ribosomal 60S subunit protein L10 |
| 10 | YOL086C | XV | 159548 | 160594 | 1,046 | 1,592 | Alcohol dehydrogenase |
| 11 | YOL039W | XV | 254297 | 254617 | 320 | 338 | Ribosomal protein P2 alpha |
| 12 | YPR080W | XVI | 700594 | 701970 | 1,376 | 1,061 | Translational elongation factor EF-1 alpha |

Two of the transcripts identified mapped to isozymes of GAPDH, forms of the same enzyme that are encoded by similar genes at different loci. The sequences of the two genes are 95.8% identical, differing at multiple variants dispersed throughout the sequences (data not shown). Even though the single-read accuracy of the direct RNA data is currently below this, further analysis of the reads mapping to each isozyme imply correct placement: firstly, multimapping (and hence randomly placed) reads are in the minority, and when these are removed, we are still left with 3217 reads mapping to YGR192C and 743 mapping to YJR009C **(Table 2)**. Secondly, there are 42 positions which differ between the two isozyme genes, and in the reads mapping to YGR192C, the most frequent nucleotide at each position was the correct base. The same was true for 42 SNPs positions in the reads mapping to YRJ009C. Thirdly, when the reads mapping to one isozyme reference sequence were analysed and variants matching that same reference were counted and plotted, alongside the variants mapping to the other isozyme reference sequence, a clear difference can be seen (**Figs. 2c and 2d**).

### ii) Direct RNA sequencing of human rhinovirus

We prepared a 1D RNA template from the ˜7.5 kb human rhinovirus (HRV) single-stranded RNA genome using the library preparation protocol outlined above and sequenced the library as before.**Fig. 3a** shows the current trace (‘squiggle’) from a full-length strand of HRV. The data was basecalled using the bespoke 1D Hidden Markov Model-based basecaller described previously, and reads were aligned to the norovirus, HRV and influenza A reference sequences using LAST. **Fig. 3b** shows the alignment, beneath a Savant representation of the NCBI Reference sequence. No off-target mapping was observed.

**Fig. 3.**
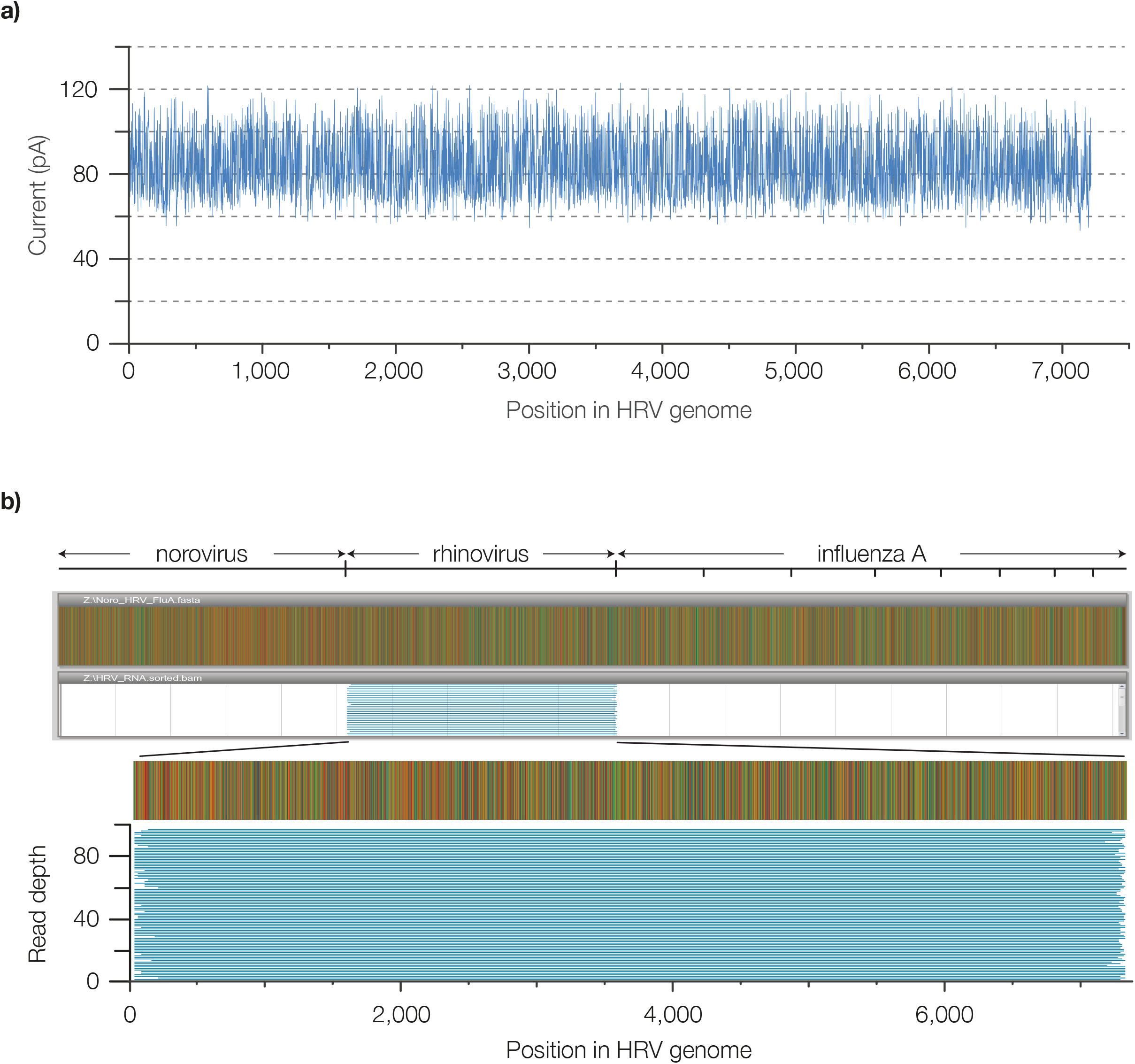
Full-length direct RNA reads of human rhinovirus. **a)** squiggle of full-length read **b)** aligned full-length reads

### iii) Direct measurement of RNA reduces bias

Having no amplification as part of the library prep means that direct RNA sequencing is free from the bias and size limitations of this step. Additionally, although reverse transcription was used in the library prep shown here, the cDNA strand served to improve performance of the system, and was not sequenced itself, so the biases of this step might be expected to have a greatly reduced influence on direct RNA sequencing compared to cDNA sequencing. To investigate this, we pooled 3 transcripts, generated synthetically from Lambda phage amplicons, in 10:30:60 ratios and counted reads of each. **Fig. 4** shows that the read counts were close to the expected values, indicating low bias.

**Fig. 4.**
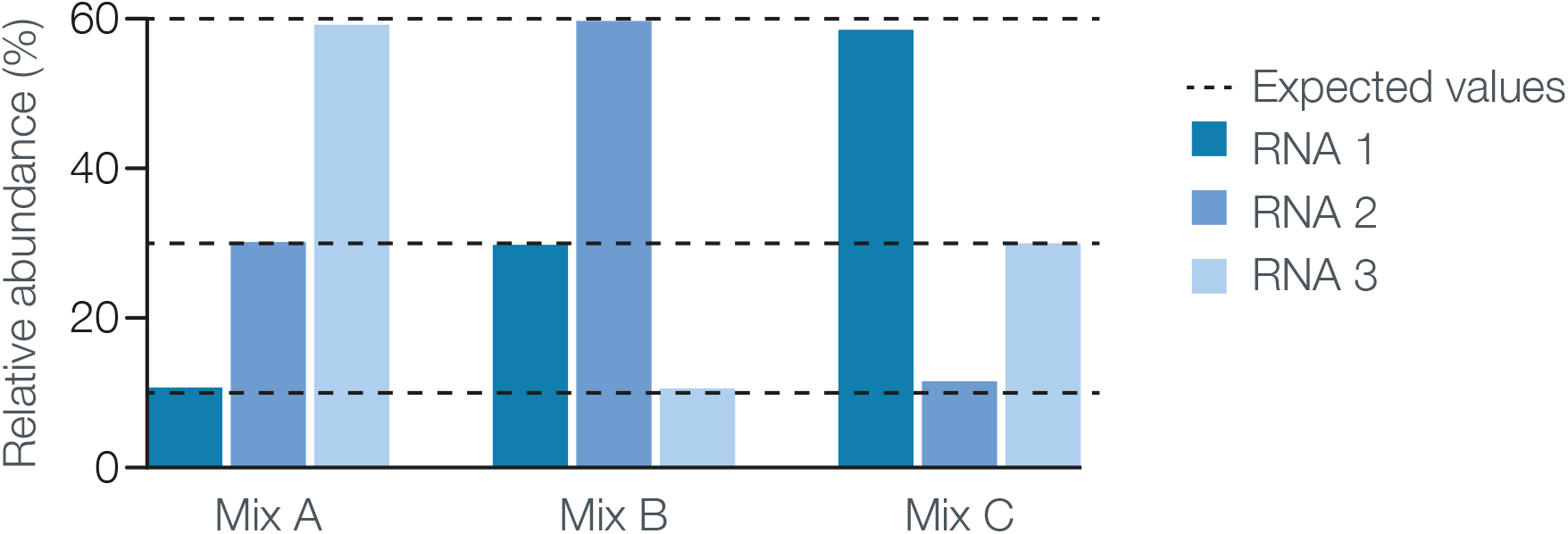
Quantitative measurement of Direct RNA reads.

### iv) Direct RNA sequencing detects modified bases

Direct RNA sequencing measures the current blockade caused by RNA bases in the narrowest part of the nanopore. Base modifications that affect this current can also be analysed. To determine the effect of a common RNA modification (m6A) on the current trace we sequenced fully modified and unmodified strands of the FLuc transcript and aligned the current traces. **Fig. 5** shows a region of this alignment. The levels corresponding to unmodified and fully modified strands are shown in red and blue respectively, beneath the base sequence of central nucleotide in the 5-mer used for alignment. m6A-containing kmers can be discriminated from rAMP-containing kmers by the nanopore.

**Fig. 5.**
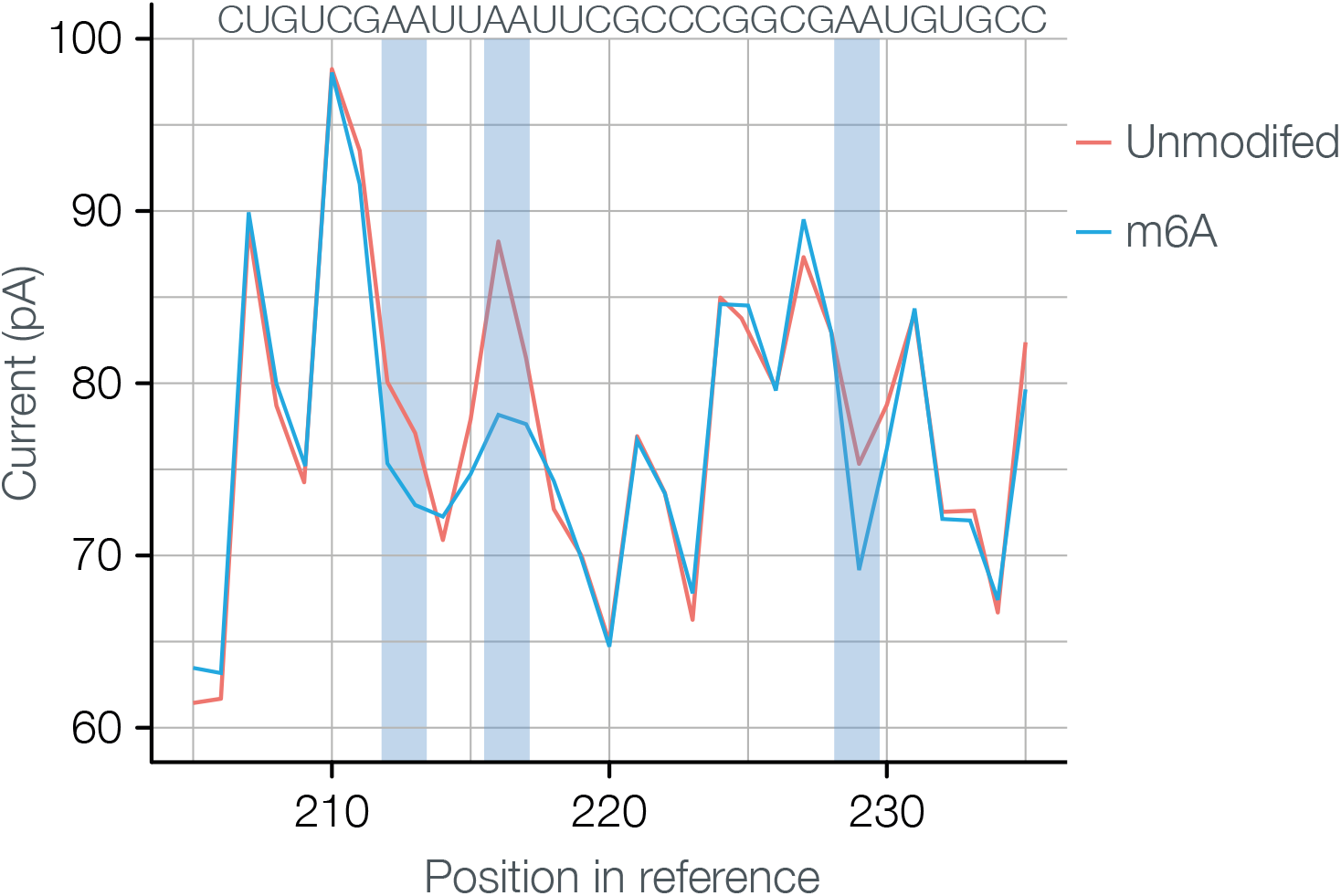
Detecting the m6A modification by direct RNA sequencing.

## Discussion

In this manuscript we present the first method for sequencing RNA in which the RNA strand is read directly, rather than by detecting the products of a synthesis reaction. This approach has many potential advantages over other RNA-seq strategies, namely that i) it is amplification-free, so should be less biased than methods which rely on either PCR or on clonal amplification, ii) by measuring the RNA directly, we avoid masking the presence of nucleotide analogues iii) our approach is compatible with very long reads, making it useful for the study of splice variants, and iv) the method is strand-specific, allowing the unambiguous identification of sense and antisense transcripts.

There is considerable room for improvement in all aspects of the method, particularly those governing throughput and accuracy. The Oxford Nanopore sequencing technology is modular, allowing straightforward modification and optimisation of all parts of the library preparation, sequencing, and data interpretation and analysis processes.

For the work presented here, we operated the motor protein at a speed of around 80 nucleotides per second. Refinements to our DNA sequencing process have allowed us to increase the DNA motor speed approximately 10-fold, with scope to increase this further. Further screening of motor protein mutants will allow us to find enzymes with the best combination of steady movement, good processivity and high processing speed increase, which will increase throughput and data quality.

The method of library preparation allows the RNA strand to pass 3’ end first through the nanopore. For DNA templates, we ordinarily sequence 5’ end first, resulting in smooth strand movement, and hence high quality data. For direct RNA sequencing, we will continue to investigate sequencing in both 3’-5’ and 5’-3’ directions.

It is not obligatory to synthesise a cDNA strand opposite the RNA template prior to sequencing, although we have found that the duplex improves throughput, possibly by reducing intramolecular secondary structure of the RNA. The library preparation method used here involved cDNA synthesis from the poly A tail of the RNA strand. This may not be ideal, because in addition to depending on high processivity of the reverse transcriptase to synthesise full-length cDNAs, it is necessary to add a poly A tail to the 3’ ends of strands which do not ordinarily have one. Future work will involve investigating other approaches to cDNA synthesis, such as random priming. However, it should be noted that because we do not sequence the cDNA strand, incomplete cDNA synthesis does not prevent sequencing of an RNA strand, and the error rate of the reverse transcriptase is not an issue.

It is possible to link the cDNA and RNA strands together with a hairpin adapter, and to sequence one strand after the other, combining the two reads into a single sequence, in an analogous way to the 2D reads generated by some Oxford Nanopore’s genomic DNA protocols. Using a motor protein that has sufficiently good movement on both RNA and DNA polynucleotides it might therefore be expected that sequencing both RNA and cDNA strands would give improved data accuracy.

For direct RNA sequencing we are currently using the same E. coli CsgG-derived nanopore that we use for DNA sequencing^24^. This has the advantage of allowing both DNA and RNA strands to be sequenced together on the same flow cell, but there may be superior nanopores for RNA sequencing. Improvements in the quality of the nanopore signal from RNA are likely to result from pore mutations and we will continue to both engineer our current pore and assess different nanopores in order to find the optimal pore for RNA sequencing. Alongside this work, we will evaluate alternative buffer compositions and running conditions, to find the best combination for high throughput and accuracy.

The basecaller used in this work is a naive HMM based solely on event means and has not been extensively optimised for the purpose of basecalling RNA data. The MinION data is far richer than this simple model supports. Logical extensions under the HMM framework would be explicit modeling of event noise or event-length distributions. It would also be beneficial to incorporate dwell time information and common signal irregularities into the model, such as periods of systematic shifts in current spanning multiple events. Improved HMMs facilitate the training of more complex models requiring pre-labeled data sets. Recurrent Neural Networks (RNNs) have been shown to be effective at basecalling MinION experiments with DNA samples^25^ and could equally be applied to RNA experiments. These models are not constrained by the Markov assumption implicit in the models used here, and can better exploit, for example, long-range correlations in the signal.

In the current work we have shown that MinION data presents a clear systematic difference between groups of molecules differing only in the presence or absence of modified nucleotides. Such analysis indicates that it is possible to detect base modifications at a single molecule level, with single-nucleotide resolution. It is trivial to extend the state space of an HMM to include states for modified bases, or to allow an RNN to provide classification scores for a larger kmer set, which will enable us to construct a basecaller to achieve this goal. Although the computational cost of HMMs will grow exponentially with the number of bases included in the model, many cases of practical relevance will wish to target limited choices of base analogues at specific loci in a reference sequence. In such cases more direct methods can be applied which need not suffer from scaling issues.

At the current level of accuracy, it will be possible to answer important biological questions, such as those involving splice variation. However, we are confident that with optimisation and further engineering of the components of the sequencing system described here we will reach a 1D accuracy and throughput for RNA sequencing that is similar to that of 1D DNA reads. We expect that the features of nanopore-based direct RNA sequencing, a combination of being amplification-free, capable of detecting nucleotide analogues, and able to generate full-length, strand-specific reads, will enable us to gain a far deeper understanding of transcriptomes than has been possible with indirect, short read sequence data.

